# From southern Africa and beyond: historical biogeography of a monocotyledonous bulbous geophyte

**DOI:** 10.1101/2022.09.21.508857

**Authors:** Cody Coyotee Howard, Leevi Nanyeni, Neduvoto Mollel, David Chuba, Alexandre R. Zuntini, Panagiota Malakasi, Timothy S. Harvey, Nico Cellinese

## Abstract

**Aim:** Within sub-Saharan Africa, plants inhabiting more seasonal and arid landscapes showcase unique distributional patterns that hint at fascinating evolutionary histories. The Ledebouriinae (Scilloideae, Asparagaceae) are widespread throughout such climates in sub-Saharan Africa, and Madagascar, the Middle East, India, and Sri Lanka. Long-distance dispersal has been hypothesized as leading to such a widespread distribution; however, low taxon sampling and taxonomic uncertainties have made uncovering the history of the Ledebouriinae difficult. Here, using the most comprehensive sampling of the lineage to date, we hypothesize that both vicariance and dispersal events impacted the biogeographical history of these bulbous monocots within and outside of Africa.

**Location:** Sub-Saharan Africa, Madagascar, Asia

**Taxon:** Ledebouriinae (Scilloideae, Asparagaceae)

**Methods:** We infer age estimates using penalized likelihood as implemented in treePL. Capitalizing on our broad geographic sampling, we use ‘BioGeoBEARS’ to reconstruct ancestral ranges and investigate the role of vicariance and dispersal.

**Results:** Our results suggest the Ledebouriinae originated within the past ∼30 myr in southeastern sub-Saharan Africa, with the major subclades arising soon thereafter. Although long-distance dispersal cannot be fully ruled out, our results lead us to hypothesize vicariance was the major process responsible for the current distribution of *Ledebouria* in Eurasia. We recover two distinct *Ledebouria* groups that overlap in eastern Africa, but are divided into mostly northern and southern clades with divergent biogeographical histories, and each showing an independent dispersal to Madagascar. A similar north-south split is seen in *Drimiopsis*. Additionally, we recover a complex biogeographic history in the predominantly sub-Saharan African *Ledebouria* clade, with a rapid radiation estimated at ∼14 mya.

**Main conclusions:** We recover evidence to suggest that the expansion of seasonal rainfall and aridity in sub-Saharan Africa, coupled with orogeny, may have fostered the diversification of the Ledebouriinae and many subclades. Miocene-driven aridification may have caused fragmentation of a once widespread distribution that led to their occurrence in Eurasia.

## 1 INTRODUCTION

Modern-day Africa is dominated by arid and semi-arid landscapes that contain a diversity of habitats from deserts, woodlands and savannas, to name a few (Bobe, 2006; Linder, 2014). In sub-Saharan Africa, these drier ecosystems collectively form a fairly continuous sickle-shaped corridor that connects the floras of southwestern, northeastern and western Africa, and that skirts around the wet tropics of central and western Africa (Balinsky, 1962; Bellstedt et al., 2012; Jürgens, 1997) (Figure 1). The gradual increase in aridification in Africa is hypothesized to have been caused by the synergistic activities of rapid global cooling, tectonic events (e.g., Eastern African rift) and oceanic upwelling (i.e., Benguela current) that altered precipitation patterns across the continent (Bobe, 2006; Couvreur et al., 2021; Hagen et al., 2021; Linder, 2017; Senut et al., 2009; Sepulchre et al., 2006). These changes had immense impacts on the evolutionary trajectories of many lineages on the African continent. For example, in the early Eocene, equatorial Africa was largely covered by tropical forests, but aridification repeatedly fragmented and reduced these habitats over time (Couvreur, 2015; Couvreur et al., 2008; Hagen et al., 2021). Evidence from plant fossils and phylogenetic studies suggest that lineages with past widespread distributions across the continent, as well as between eastern Africa and southern Asia, are now disjunct as a result of recent aridification (Ali et al., 2013; Jacobs et al., 1999; Pokorny et al., 2015; Sanmartín et al., 2010; Zhou et al., 2011). Conversely, dry and more seasonal climates have been linked to an increase in the diversification and dispersal of lineages both in and out of Africa, and today, a rich diversity of taxa can be found within such climates in Africa as well as surrounding areas (e.g., Arabian Peninsula) (Bruyns et al., 2014; Coe & Skinner, 1993; Jürgens, 1997; Lorenzen et al., 2012; Nylinder et al., 2016). Therefore, it is imperative we gain a broad understanding of the historical evolution of diverse lineages across different landscapes, habitats, and biomes, especially in fragmented regions, in order to continually refine our understanding of the past and improve our ability to predict the ways in which African biodiversity may be shaped in the future.

**Figure 1.**
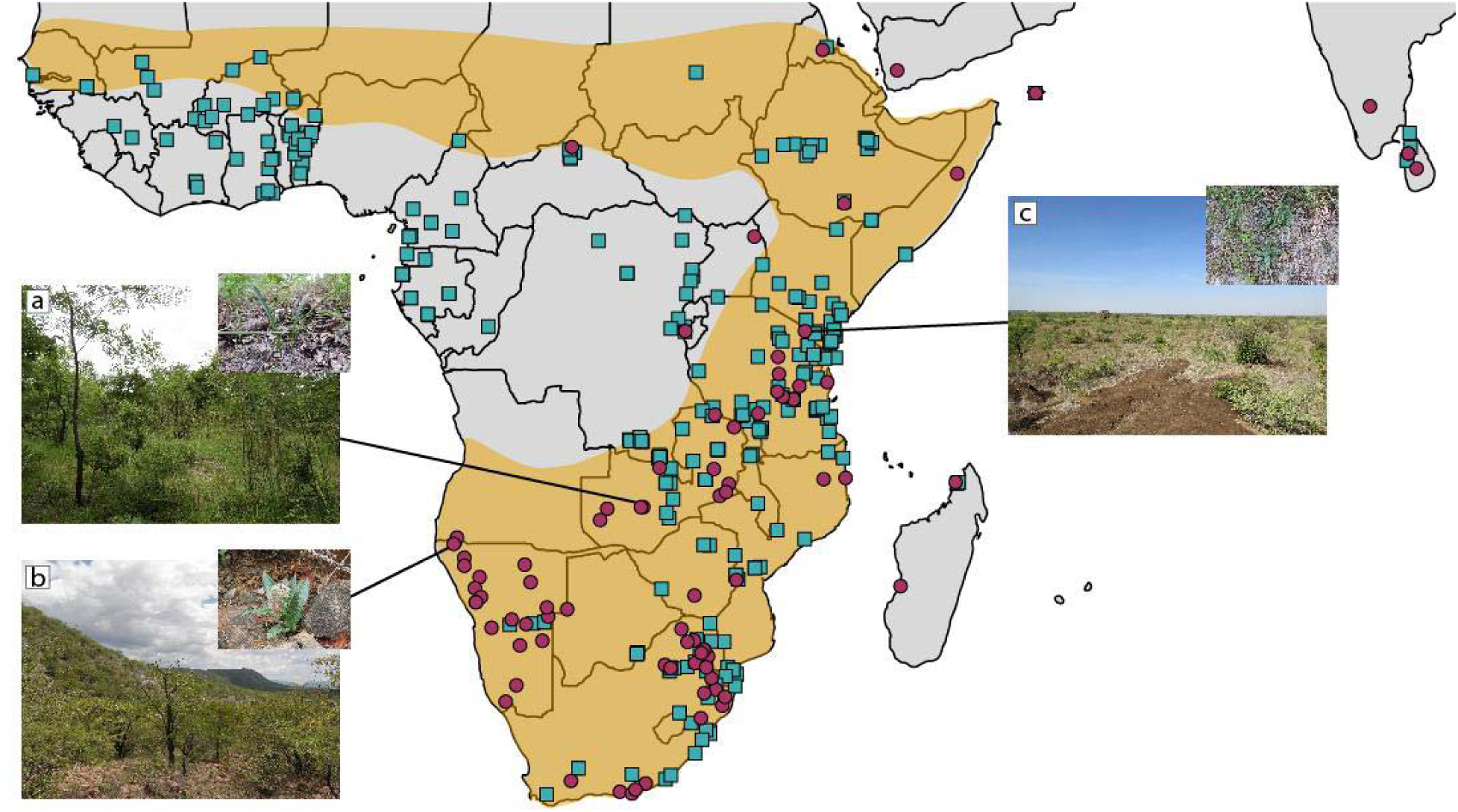
General distribution of the seasonal and arid ecosystems of sub-Saharan Africa (light orange polygon) underlain by the general distribution of the Ledebouriinae, which is sufficiently captured by displaying GBIF specimen occurrence data (teal squares; GBIF.org (22 March 2022) GBIF Occurrence Download https://doi.org/10.15468/dl.xmccae) and the location of samples collected for this study (maroon circles). Do note though that sampling gaps exist, for example in Angola, Nigeria, and South Sudan. Examples of habitats and associated Ledebouriinae taxa are displayed in the images. a) *Ledebouria* sp. 8 CCH186, b) *Ledebouria* sp. 15 CCH027, c) *Drimiopsis botryoides* subsp. *botryoides* CCH153. See Figure 2 for phylogenetic placement of these example taxa. General distribution of seasonal landscapes adapted from Balinsky (1962), Jürgens (1997), and Bellstedt et al. (2012).

Geophytes, herbaceous plants with renewal buds located belowground on structures such as bulbs, corms, and stem tubers, are ubiquitous components of seasonal or disturbance-prone habitats, and are phylogenetically diverse (Howard et al., 2019; Pausas et al., 2018; Tribble et al., 2021). Within Africa, geophytes are major components of the Greater Cape Floristic Region and the Mediterranean Basin (Buerki et al., 2012; Procheş et al., 2006). Although geophytes are predominant elements of these two areas, the geophytic habit is widespread throughout sub-Saharan Africa, particularly within seasonal or disturbance-prone (e.g., fire) habitats (Esler et al., 1999; Kornas, 1985). Studies have reported that many African geophytes’ origins coincide with the onset of increased seasonality and/or aridity within the continent (i.e., since the Eocene), with the majority of diversity evolving in response to the relatively more drastic climatic changes since the Oligocene/Miocene (Ali et al., 2012, 2013; Buerki et al., 2012; del Hoyo et al., 2009; Procheş et al., 2006). At a broad scale, therefore, geophytic lineages make excellent candidates for understanding relatively recent evolutionary and biogeographic dynamics within seasonal and arid climates.

Scilloideae (Asparagaceae) are a bulbous geophyte lineage widespread both within and outside of Africa (Speta, 1998a). This monocotyledonous clade consist of 1,000+ taxa found throughout seasonal climates in Africa as well as Madagascar, Europe, the Middle East, and Asia, with a single lineage found in South America (i.e., *Oziroe* Raf.; Giranje & Nandikar, 2016; Speta, 1998a). Although only a handful of studies have investigated the historical biogeography of the Scilloideae and its subclades, so far, all have pointed to sub-Saharan Africa as the origin for the majority of the group (excluding *Oziroe* in South America), followed by a complex history within and outside of the continent (Ali et al., 2012, 2013; Buerki et al., 2012; Pfosser, 2012). Age estimates for the Scilloideae suggest an origin within the past ∼70–50 myr, and major subclades within the past ∼60–50 myr (Ali et al., 2013; Buerki et al., 2012); however, most inferences have been made with low taxon sampling, and in biogeographical studies sub-Saharan Africa has been broadly subdivided into one to three areas (Ali et al., 2012, 2013; Buerki et al., 2012; Pfosser, 2012). Regardless, given the current age estimates, we hypothesize that the majority of Scilloideae lineages diversified in response to an increase in climatic seasonality and aridification that intensified from the Eocene onwards. Focusing on widespread groups within the Scilloideae using a more detailed approach to the regionalization of Africa as well as a greater taxon sampling may provide refined insights into the biogeographical processes that have impacted the dispersal of the Scilloideae as well as other plant lineages found across and outside of Africa.

The Ledebouriinae are an ideal group to study because they are widespread within sub-Saharan Africa, with a handful of taxa found in Madagascar, Socotra, Yemen, India, and Sri Lanka (Giranje & Nandikar, 2016; Venter, 1993) (Figure 1). This distribution is unique within Scilloideae since many sympatric lineages with Ledebouriinae are also found in northern Africa (Pfosser, 2012; Speta, 1998b), but the Ledebouriinae are absent from the Scilloideae-rich Mediterranean Basin (Venter, 1993, 2008) (Figure 1). In sub-Saharan Africa, the Ledebouriinae are predominantly found within regions that experience oscillating weather patterns of wet and dry and/or cold and warm, with highest diversity in the Limpopo, Mpumalanga, and KwaZulu-Natal regions of South Africa (Venter, 1993), yet some occurrences are documented from more wet, tropical regions (e.g., Gabon) (Figure 1). Much diversity within the Ledebouriinae, however, remains undescribed to science (Howard, 2014; Howard et al., 2022). Additionally, an expanded phylogenomic analysis of the group suggests that a complex biogeographical history awaits to be thoroughly examined (Howard et al., 2022). Previous dating analyses have estimated the origin of the Ledebouriinae sometime within the last ∼25 myr in sub-Saharan Africa (Ali et al., 2012; Buerki et al., 2012). However, this clade was not the focus of study and therefore, taxon sampling was low. Additionally, results were unable to provide fine-scale biogeographical patterns since sub-Saharan Africa was considered and analyzed as one large area. Regardless, given our current understanding of the Ledebouriinae distribution, their dispersal ability, and a recent phylogenomic study on this group (Howard et al., 2022), we can generate several hypotheses regarding major biogeographical events throughout their history.

Although Ledebouriinae diversity is highest within Africa, many taxa are also found outside of the continent (Venter, 1993). However, this intriguing distribution has yet to be the explicit focus of investigation. Previous phylogenetic studies have hypothesized that multiple dispersals to Madagascar from mainland Africa have occurred within the Ledebouriinae (Pfosser, 2012). For example, within Madagascar, we find two endemics: *L. nossibeensis* and an undescribed taxon with morphological affinities to *L. socialis* (Pfosser, 2012). In addition to their morphological differences, these two Malagasy taxa are also found in distinct regions of the island (northwest vs southwest; see Figure 1). Madagascar has also been hypothesized to have played a role in the dispersal of *Ledebouria* to India. For example, the phylogenetic reconstructions of Ali et al. (2012) recovered Malagasy and Indian *Ledebouria* as sisters, which led the authors to invoke long-distance dispersal from Madagascar to India. However, as stated by the authors, taxa from eastern, western, and northern sub-Saharan Africa were absent, limiting confidence in these conclusions (Ali et al., 2012). Furthermore, low phylogenetic resolution from Pfosser (2012) diminishes confidence in the hypothesized independent dispersal events to Madagascar. Recently, a phylogenomic analysis of the Ledebouriinae provided the framework to suggest (without testing) multiple dispersals to Madagascar, and a potential migration out of Africa via the Arabian Peninsula into India (Howard et al., 2022). Therefore, we hypothesize that at least two independent dispersals to Madagascar have occurred in the Ledebouriinae. This is based on the distinct distribution and morphotypes of Malagasy taxa, as well as their independent phylogenetic placement (Howard et al., 2022; Pfosser, 2012). Regarding dispersals out of Africa, we hypothesize that a once widespread distribution between Africa and India by way of the Middle East fragmented due to Miocene-driven aridification, isolating populations and contracting the overall distribution ranges. This hypothesis is drawn from several lines of evidence. First, *Ledebouria* seed has limited dispersal capacity, which occurs primarily via sheet water flow (i.e., dispersal occurs only short distances and is rainfall dependent) (Venter, 1993). In years of limited rainfall and/or in flat terrains, seeds and seedlings can be found surrounding the parent plant (CCH, pers. obs.), and the ephemeral nature of the seeds in some taxa warrants germination soon after fruit dehiscence (CCH and TSH, pers. obs.). Therefore, we hypothesize that long distance dispersal from Africa or Madagascar to India is unlikely. Further evidence for a once-widespread, now-fragmented distribution is found in the current distribution of the Ledebouriinae in the Arabian Peninsula. Today, *L*. *yemenensis* is the only Ledebouriinae currently recorded from the region, and it is endemic to the cooler, higher elevations of the Yemeni Highlands that are surrounded by arid lowlands. We hypothesize this taxon represents a relic of a once much larger range. Additionally, there are multiple, endemic taxa to the Socotra archipelago (Miller & Morris, 2004). For example, *L. grandifolia* and a putative *L.* aff. *revoluta* are found only on the island of Socotra, and *L. insularis* is endemic to the summit of Samha, where low clouds provide a moist habitat relative to the surrounding desert landscape (Miller & Morris, 2004). Therefore, given the multiple taxa located on different islands in the Socotra archipelago, we hypothesize that vicariance led to allopatric speciation as the islands moved farther away from the mainland. In summary, although the Ledebouriinae occur within seasonally dry and arid regions, many taxa may not be adapted to intense aridity, such as that found in much of the Middle East and Pakistan, that intensified during the Miocene.

The unique, widespread distribution of the Ledebouriinae provides us with the opportunity to refine our understanding of evolution and biogeography within Africa, potentially during a time of extensive climatic, geologic, and habitat change on the continent. Here, using a detailed categorization of the African continent, we investigate the timing and historical dispersal of the Ledebouriinae both within and out of Africa to test major biogeographical hypotheses: 1) due to high diversity within southern Africa, this region may represent the putative origin of the Ledebouriinae; 2) at least two independent dispersals occurred to Madagascar from mainland Africa; 3) vicariance is the major biogeographical process leading to the widespread distribution of *Ledebouria* in Eurasia. Our increased sampling of the Ledebouriinae allows for a more thorough investigation into the evolutionary history of this group and will improve our knowledge of biogeographical patterns and processes within Africa.

## 2 METHODS

### 2.1 Phylogenetic analysis

Ledebouriinae samples were obtained from the field, private collections, and herbarium vouchers (Howard et al., 2022). DNA extractions were performed using a modified CTAB protocol, followed by high throughput sequencing on an Illumina HiSeq using the Angiosperms353 universal probe set (Johnson et al., 2019). Raw reads were cleaned using SECAPR (Andermann et al., 2018), sequences were pulled using HYBPIPER (Johnson et al., 2016) and aligned using MAFFT v.7 (Katoh & Standley, 2013). See Howard et al. (2022) for more details on Ledebouriinae sequence acquisition and analysis as well as data used, which can all be found on Dryad (https://doi.org/10.5061/dryad.nzs7h44q6).

We incorporated outgroup taxa from the Plant and Fungal Tree of Life project (Baker et al., 2022) and the 1KP dataset (Matasci et al., 2014). Exons were only available for the outgroup taxa included to estimate divergence times using fossils and secondary calibration points. Phylogenetic reconstruction including the Ledebouriinae plus outgroups was performed on a concatenated, partitioned supermatrix of exons with 10% gaps/ambiguous sites removed using phyx (Brown et al., 2017). This matrix was analyzed using IQ-TREE v.2-rc1 (Nguyen et al., 2015) with 1000 ultrafast bootstraps, a best-fit partitioning scheme using the greedy algorithm of PARTITIONFINDER (Lanfear et al., 2012), and a relaxed clustering percentage of 10 (Lanfear et al., 2014), followed by phylogenetic reconstruction (-m TESTMERGE). However, the exon-only dataset returned low support for many nodes within the Ledebouriinae. Therefore, we reran the IQ-TREE analysis with a topological constraint tree (-g) that was previously built from a supercontig (i.e., exons + introns) dataset of the Ledebouriinae (Howard et al., 2022).

### 2.2 Time calibration

We incorporated eight outgroup fossil calibration points, each with a minimum age specified in Iles et al. (2015) (Table S1). A secondary calibration point at the crown node of monocots was inferred between 131–135 mya based on previous analyses (Givnish et al., 2018; Magallón et al., 2015).

Given the size of the dataset, we used penalized likelihood as implemented in TREEPL (Smith & O’Meara, 2012) for time calibration. To incorporate uncertainty around age estimation, we took a multi-tiered approach (see https://github.com/sunray1/treepl) similar to previous studies estimating divergence times using large phylogenetic datasets (Emberts et al., 2020; Li et al., 2019; Magallón et al., 2015). We generated 100 bootstrap replicates of our original exon supermatrix alignment using RAXML v.8.2.0 (f-j option) (Stamatakis, 2014). A maximum likelihood tree for each replicate with a corresponding partition file was then reconstructed using a topological constraint (i.e., the phylogeny from the IQ-TREE analysis) to ensure consistent calibration point placement, and a GTRGAMMA model of evolution. The resulting 100 “best trees” were rooted on *Acorus gramineus* using phyx (Brown et al., 2017). In Step 1, each replicate tree had a priming step completed with a random seed number and the thorough command invoked. Step 2 was performed three independent times to assess convergence on the best smoothing parameter for each individual tree. Step 2 also included the individual outputs from each tree’s previous priming step (e.g., optad, moredetail, etc.) as well as the cross-validation (CV) steps, which were set to cvstart=10000, cvstop= .00000000000001, cvmultstep=0.09. Lastly, Step 3 summarized each individual tree’s CV output to determine and scale each tree using the appropriate smoothing parameter. The 100 ultrametric phylogenies were summarized using TREEANNOTATOR v.1.10.4 (Bouckaert et al., 2014) to obtain a maximum clade credibility tree with 95% confidence intervals around each node and median node heights.

### 2.3 Biogeographical analysis

The combined biogeographical regionalization of Linder et al. (2012) (see Figure 2 in Linder et al., 2012) was used for categorizing the location of each sub-Saharan African Ledebouriinae accession. The exact distribution of populations/taxa represented by many field-collected individuals (i.e., collections made by C.C. Howard) remains to be fully assessed since they are undescribed species and/or are only known from one locality. Additionally, many described species are currently known as occurring in small geographic ranges or even single mountain tops (Lebatha, 2004; Venter, 2008). Consequently, most samples were assigned only to known areas of occurrence or areas they were collected. Likewise, accessions found outside of Africa were coded according to their actual known provenance: Madagascar, Yemen, Socotra, India, or Sri Lanka. Even when seemingly widespread, taxa were coded using their documented occurrence, rather than their putative distribution inferred by ambiguous taxonomic knowledge. For example, *Ledebouria revoluta* is reported from the Cape Region of South Africa to Sri Lanka (Mwafongo et al., 2017; Venter, 1993), however, this widespread taxon is poorly understood, and it may harbor undescribed diversity as evidenced by the wide range of morphological variation across its entire distribution range (Stedje, 1998). Its diversity has not been investigated and therefore, this name is often applied (or misapplied) to individuals whose species identification is difficult (Mwafongo et al., 2017). Consequently, and not surprisingly, *L. revoluta* has become a ‘bucket’ species, an ambiguous and complex entity as evidenced by its polyphyly (Howard et al., 2022).

**Figure 2.**
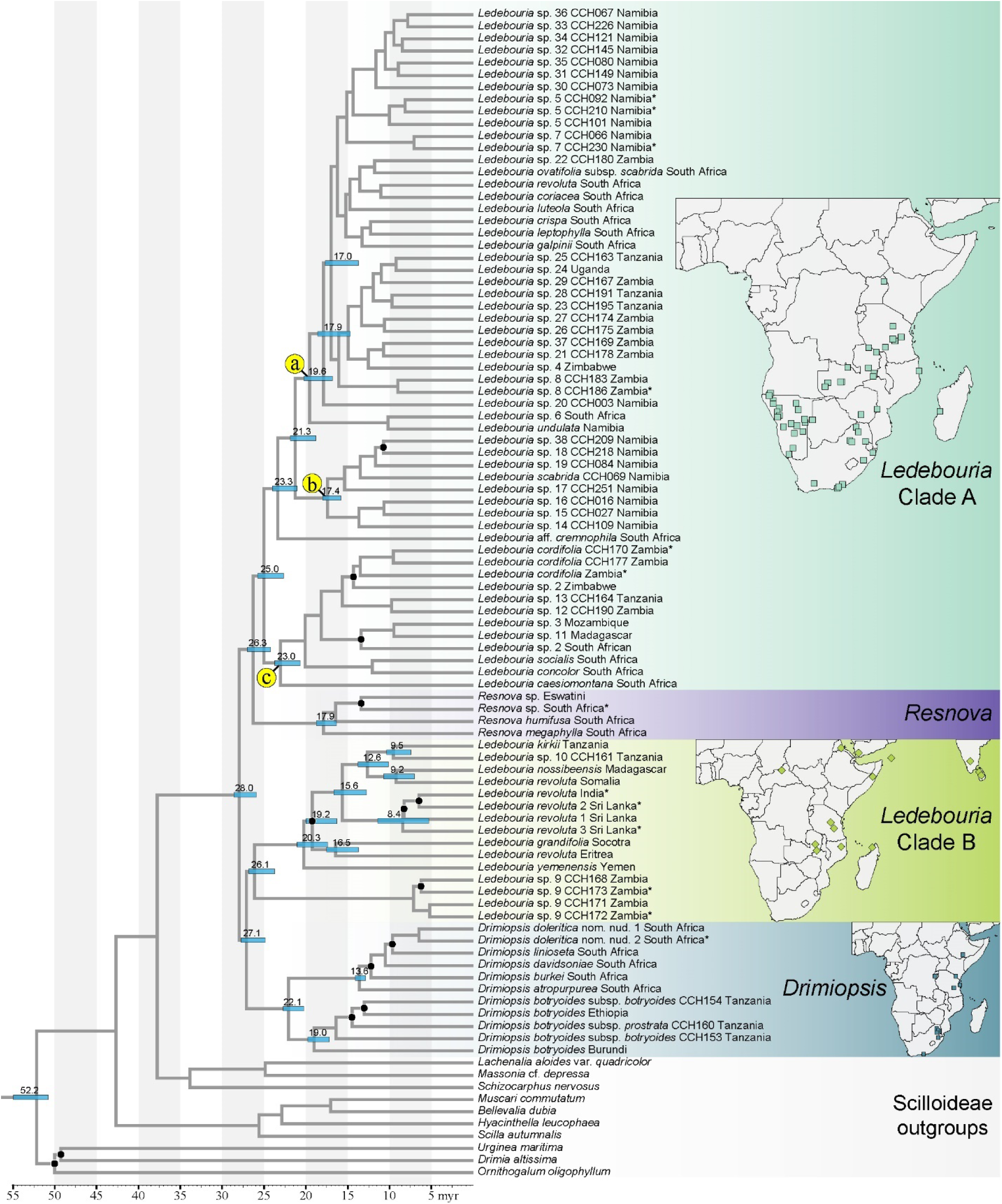
Divergence times within the Ledebouriinae as estimated using penalized likelihood in treePL. For each clade, collection localities are overlaid on a map of Africa, Madagascar, and Asia to aid with discussions of geographical patterns. Clades labeled with a, b, and c in yellow are clades discussed within the manuscript. Black circles indicate nodes with ultrafast bootstrap support values below 90 as recovered from the IQ-TREE analysis. Numbers above blue bars denote median age estimates. Blue bars denote 95% confidence intervals. * indicates samples excluded from subsequent biogeographical analyses. Not all nodes are annotated with estimated divergences times for illustrative simplicity. For a fully annotated phylogeny, see Figure S1.

In total, 14 geographic regions were used to code taxa. To estimate ancestral ranges, we compared the maximum likelihood implementations of three biogeographical models, DEC, DIVA-like, and BAYAREA-like as implemented in ‘BioGeoBEARS’ (Matzke, 2013a, 2013b, 2014) in R (2016). We used log-likelihood and AICc values to determine the best model between the three. Given the debate surrounding the use of models with the +j parameter (Matzke, 2022; Ree & Sanmartín, 2018), we decided to not include them and, additionally, their implementation tends to infer a high number of jump dispersals, which are unlikely biological scenarios for the majority of the Ledebouriinae (see Introduction and Discussion).

#### 2.3.1 Biogeographical uncertainty

Within a subclade of *Ledebouria* Clade A, phylogenetic relationships are poorly supported (Figure 2, node a) (Howard et al., 2022). However, given the potential rapid radiation along the backbone of this group, we wanted to explore the biogeography of this clade since it may contain an interesting history. To compensate for the uncertainty, we incorporated topological and branch depth variability into our biogeographical analysis by using a random sampling of trees from the posterior of a Bayesian analysis for the subclade of interest (Ceccarelli et al., 2019; Magalhaes et al., 2021). To begin, we reduced our total alignment to the most clock-like genes using scripts available in SortaDate (Smith et al., 2018). This was done due to failure to reach convergence when using the entire alignment (data not shown). We kept genes that were at least 10% concordant (bipartition > 0.1), had a tree length greater than 7.24, and had root-to-tip variation of less than 0.009. Cutoff values for the latter two were determined using the median values for all gene trees, as previously reported (Pillon et al., 2021). Of the remaining alignments, we included only taxa within the subclade of interest (Figure 2b) and removed duplicate Ledebouriinae outgroup taxa as well as those putatively duplicate within the Ledebouriinae ingroup (indicated by * in Figure 2). Each gene was then aligned using MAFFT invoking the--auto option. Sites with >5% gaps/ambiguity were removed using phyx. Due to failure to reach convergence using a partitioned, concatenated alignment in BEAST v1.10.4 (Drummond & Rambaut, 2007), we instead input an unpartitioned, concatenated alignment using a GTR+GAMMA model of substitution, a speciation birth-death prior, and a lognormal distribution with an offset 17.1, mean 1.0, stdev 1.0 (95% range 17.24–19.88) on the root node of the subclade. This initial value was chosen based on the treePL divergence time estimates for this node. We ran an MCMC chain of 300k generations sampling every 10k generations. Convergence was assessed using TRACER v1.7.1 (Rambaut et al., 2018) to ensure all ESS values were above 200. We removed 50% of trees as burnin and used the remaining trees in the analysis. We randomly sampled 100 trees after burnin, and performed a DIVA-like analysis (i.e., the best model) on each tree followed by stochastic mapping using ‘BioGeoBEARS’. Trees were summarized to obtain an average number of biogeographical events within and between each region included in the analysis. Scripts for running the analysis can be found on github (https://github.com/ivanlfm/BGB_BSM_multiple_trees). The rationale for incorporating phylogenetic uncertainty was developed by Ceccarelli et al. (2019) and Magalhaes et al. (2021).

## 3 RESULTS

### 3.1 Phylogenetic relationships and age estimates

We recovered a polyphyletic *Ledebouria*, and a monophyletic *Drimiopsis* and *Resnova*, which is congruent with a recent study (Howard et al., 2022). However, shallow-level relationships differed compared to Howard et al. (2022), particularly within the *Ledebouria* clades, and especially within *Ledebouria* Clade A. This is due to using a topological constraint (-g), which searches tree parameter space, over using a fixed topology (-te) which does not perform a tree search (see http://www.iqtree.org/doc/Command-Reference). Given differences in the inputs between Howard et al. (2022) and our exons-only datasets, we preferred a topological constraint allowing for a tree search over forcing a fixed topology. One major difference between our results and those of Howard et al. (2022) can be found in *Ledebouria* Clade B, specifically the subclade containing mostly non-African taxa (i.e., *L. grandifolia* + *L. revoluta* India). Howard et al. (2022) used supercontigs (exons + flanking intronic region) for phylogenetic reconstruction, and they recovered *L. grandifolia* (Socotra) as sister to a clade containing *L. yemenensis* (Yemen), which was sister to the remaining taxa. All branches within the clade were recovered with high ultrafast bootstrap support (UF-BOOT). In this study (Figure 2), using only exons, we recovered *L. yemenensis* as sister to the same clade with high support, not *L. grandifolia*, which was instead recovered as sister to *L. revoluta* (Eritrea). Additionally, the node proceeding *L. yemenensis* was recovered with low support (UF-BOOT of 76).

Within a penalized likelihood framework, the Scilloideae originated approximately 52.0 mya (95% CI 51.2–54.5) (Figure 2; Table 1). We recovered a median crown age estimate for the Ledebouriinae of 28.0 myr (95% CI 26.1–28.3), with age estimates of the four major Ledebouriinae clades soon thereafter. The split between *Drimiopsis* and *Ledebouria* Clade B was estimated at 27.1 mya (95% CI 25.2–27.5). The crown of *Drimiopsis* was dated to 22.1 mya (95% CI 20.5–23.5), and *Ledebouria* Clade B was dated at 26.1 mya (95% CI 24.0–26.5). The split between *Resnova* and *Ledebouria* Clade A occurred at 26.3 mya (95% CI 23.7–25.2). The *Resnova* crown was dated at 17.9 mya (95% CI 16.6–18.5), and *Ledebouria* Clade A was estimated at 25.0 mya (95% CI 23.0–25.4) (Table 1).

**Table 1.** Comparison of median age estimates and associated uncertainty (95% confidence intervals (CI)) for each major Ledebouriinae clade using penalized likelihood as implemented in treePL.

| Clade | median estimate | 95% CI |
| --- | --- | --- |
| Scilloideae | 52.2 | 51.2–54.5 |
| Ledebouriinae | 28.0 | 26.1–28.3 |
| *Ledebouria* Clade A | 25.0 | 23.0–25.4 |
| *Resnova* | 17.9 | 16.6–18.5 |
| Clade A + *Resnova* | 26.3 | 24.5–26.7 |
| *Drimiopsis* | 22.1 | 20.5–23.5 |
| *Ledebouria* Clade B | 26.1 | 24.0–26.5 |
| Clade B + *Drimiopsis* | 27.1 | 25.2–27.5 |

### 3.2 Biogeography

The DIVA-like model produced the most likely ancestral range estimates among the models (LnL-220.54) (Table 2). The regions Natal+Zambezian were reconstructed as the ancestral range for the Ledebouriinae (*p* = 0.53) (the next best area being Kalahari + Zambezian; *p* = 0.29) and *Drimopsis* (*p* = 0.84), while Zambezian was reconstructed for *Drimiopsis* + *Ledebouria* Clade B (*p* = 0.95) and *Ledebouria* Clade B (*p* = 0.98). Within *Ledebouria* Clade B (i.e., *L. yemenensis* + *L. revoluta* Somalia), the model strongly favors a widespread distribution of Yemen+Zambezian (*p* = 0.55) (the next best area being Zambezian; *p* = 0.38) followed by subsequent reconstructions containing various combinations of Zambezian and other regions, or Zambezian alone (Figures 3 and S2). Within *Ledebouria* Clade B we recover one of two dispersals to Madagascar (Figure 3).

**Table 2.** Statistical outputs for each model from BioGeoBEARS. log-likelihood (LnL); rate of dispersal (d); rate of extinction (e); Akaike’s Information Criterion (AIC)

| Model | LnL | Number of parameters | $d$ | $e$ | AIC |
| --- | --- | --- | --- | --- | --- |
| DEC | -267.1 | 2 | 0.01 | 0.01 | 538.4 |
| DIVA-like | **-220.5** | **2** | **0.0027** | **0.009** | **445.2** |
| BAYAREA-like | -232.8 | 2 | 0.0036 | 0.04 | 469.8 |

**Figure 3.**
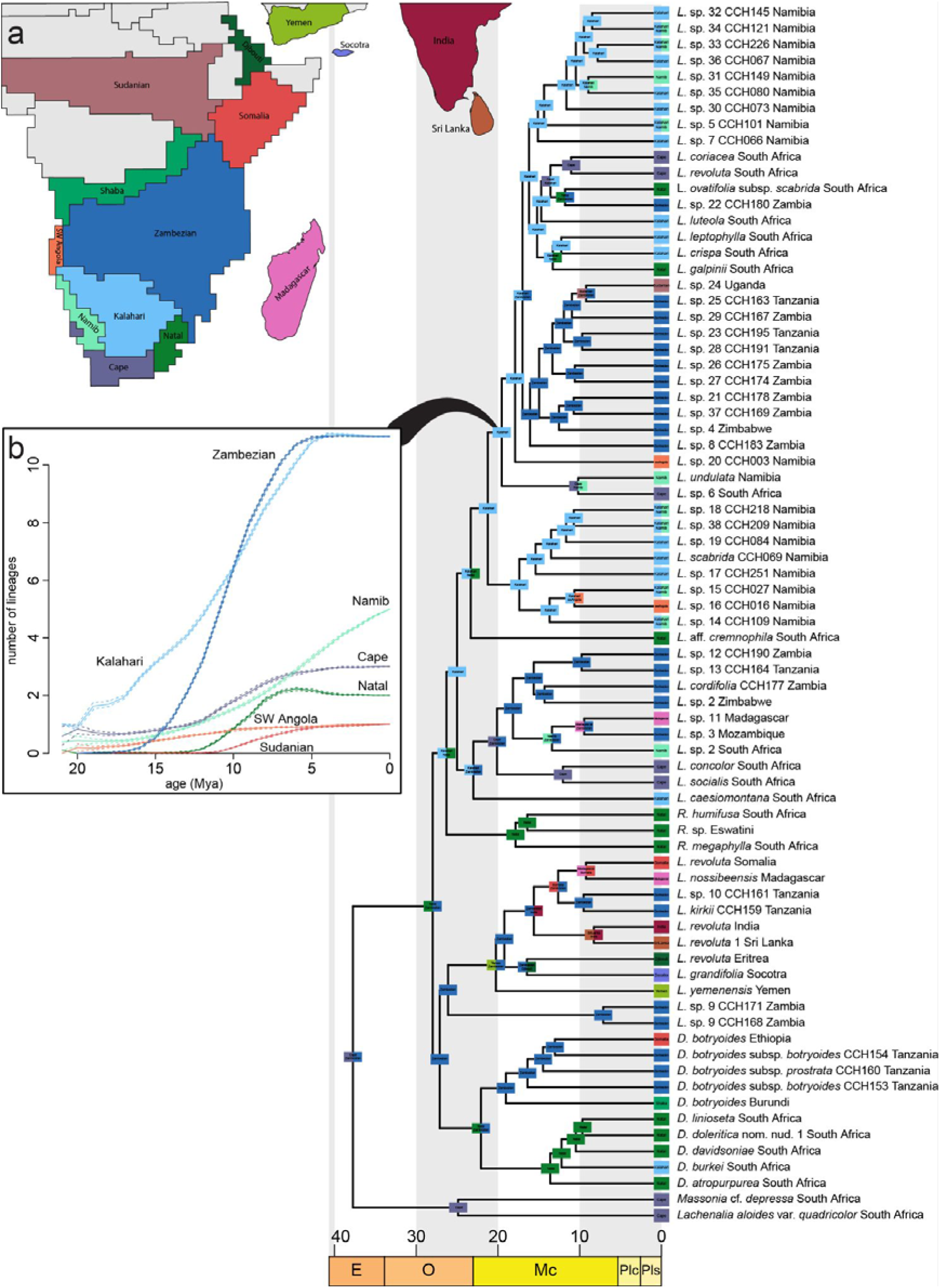
Biogeographical reconstruction using the DIVA-like model as implemented in ‘BioGeoBEARS’ and plotted using ‘RevGadgets’. Colored boxes at nodes display the region reconstructed with the highest probability; colors used at nodes match those in the upper left corner map (i.e., Fig. 3b). a) Regionalization scheme from Linder et al. (2012). used in the classification of taxa for biogeographical analysis. b) Summary of 10 stochastic mappings from a posterior distribution of trees including only taxa from the indicated clade (solid black line). For pie chart probabilities at each node, see Figure S2.

The Kalahari+Natal (*p* = 0.42) or Natal (*p* = 0.35) regions were recovered as the ancestral range for *Resnova* + *Ledebouria* Clade A, whereas the Natal region was recovered for *Resnova* (*p* = 0.99), and Kalahari for *Ledebouria* Clade A (*p* = 0.51) (the next best area being Natal; *p* = 0.18) (Figures 3 and S2). Within *Ledebouria* Clade A, we find an additional dispersal to Madagascar (Figure 3). Additionally, in summarizing the 10 biogeographic stochastic reconstructions (see section 2.3.1), we recovered a steady rise in the number of lineages in the Zambezian and Kalahari regions from ∼15 myr onwards in a subclade of *Ledebouria* Clade A (Figure 3b). Additionally, despite some differences in topology between these results and those of Howard et al. (2022), both studies overall share clades that largely reflect geography. We recover two subclades of mostly Namibian taxa, a subclade containing mostly South African taxa, and a subclade containing a mixture of Zambian, Tanzanian, and Zimbabwean taxa (Figures 2 and 3).

## 4 DISCUSSION

### 4.1 Broad biogeographical patterns

Here, we present the best and most comprehensive sampling of the Ledebouriinae to date (representing ∼30% of described Ledebouriinae taxa (POWO, 2019) plus numerous undescribed taxa), which has provided greater insights into the historical biogeography of this widespread, bulbous lineage than previously uncovered. Our results suggest a rapid radiation along the backbone of the Ledebouriinae, estimated between ∼28–26 mya (Figure 2; Table 1) in southeastern sub-Saharan Africa (Figures 3, S1). This region corresponds somewhat to the current center of diversity of the Ledebouriinae, which is at the intersection of the Natal-Kalahari-Zambezian biogeographical regions (Lebatha, 2004; Manning, 2020; Venter, 1993). The four major subclades (*Ledebouria* Clade A, *Ledebouria* Clade B, *Resnova*, and *Drimiopsis*) originated in neighboring regions to one another soon thereafter (Table 1; Figure 3). During this timeframe in Africa, major shifts in climate and habitat composition have been inferred at both local and continental scales, and at shallow and deep phylogenetic scales (Couvreur et al., 2021; Gizaw et al., 2021; Hagen et al., 2021; Kandziora et al., 2022; Pokorny et al., 2015). From the Eocene-Oligocene transition, global cooling promoted the expansion of seasonal and arid climates as well as savanna, grassland, and fire-prone habitats in Africa, with greater increases during the Miocene (Senut et al., 2009; Sepulchre et al., 2006). Furthermore, the Great Escarpment underwent renewed and sporadic uplift during the Miocene, which increased habitat heterogeneity that was coupled with an intensification of global cooling and aridification that collectively led to lineage radiation and dispersal in many African taxa (Cowling et al., 2009; Galley et al., 2007; Neumann & Bamford, 2015; Partridge & Maud, 1987). Being bulbous geophytes, which overall are found in some of the driest and most seasonal climates (Howard et al., 2019), the Ledebouriinae may have been “pre-adapted” for these climatic changes, and therefore diversified in response (Howard et al., 2022). Overall, our biogeographic reconstructions and divergence time estimations lead us to hypothesize that the expansion of seasonal habitats coupled with orogenic activity in southern sub-Saharan Africa spurred the radiation of the Ledebouriinae and promoted additional radiations and dispersals over time.

### 4.2 A tale of two *Ledebouria*

We estimate that the two *Ledebouria* lineages originated at slightly different times (Figure 2; Table 1) in neighboring regions (i.e., Kalahari vs Zambezian), which led to subsequent divergent biogeographic histories (Figure 3). In general, the two *Ledebouria* overlap in eastern Africa with a geographical divide between southern and northern sub-Saharan Africa (Figure 2) (Howard et al., 2022). The divergent evolutionary histories and current overlapping distribution of the two *Ledebouria* may have been influenced by the expansion and contraction of wet, tropical, forested habitats in eastern Africa that repeatedly split a historically more widespread distribution (Couvreur, 2015; Couvreur et al., 2021). These past habitat fluctuations have created a mosaic of climates and vegetation types in eastern Africa, and have influenced the evolutionary trajectory of many lineages (Dagallier et al., 2020; Lorenzen et al., 2012). Other lineages adapted to seasonal and arid habitats share similar distributions with the two *Ledebouria* (Grace et al., 2015; Jürgens, 1997). For example, the two lineages overlap geographically with that of stapeliads (Ceropegieae, Apocynaceae)—*Ledebouria* Clade B overlaps with the Northern grade, whereas *Ledebouria* Clade A overlaps with the Pan-African clade (see Fig. 3 in Bruyns et al., (2014)). We also find two dispersals of *Ledebouria* to Madagascar (Figure 3), which was previously hypothesized (Pfosser, 2012). Madagascar and mainland Africa have been separated since at least the Paleocene (Couvreur et al., 2021), and our results therefore suggests long distance dispersal from Africa to Madagascar led to the origin of Ledebouriinae on the island. Interestingly, both independent dispersal events are estimated to have occurred at similar times (i.e., the confidence intervals overlap) (Figures 2, 3, S1), but this timing does not match with the direction of ocean current flows that would have facilitated transport via rafts from Africa to Madagascar (J. R. Ali & Huber, 2010). However, dispersal rates for animals were estimated to not have changed due to ocean current flow direction, but instead may have increased after the current shifted to an East to West flow direction (Samonds et al., 2012). Although completely speculative, but interesting to think about nonetheless, perhaps an environmental phenomenon, such as a historical major cyclone (Samonds et al., 2012), facilitated overwater dispersal of Ledebouriinae from Africa to Madagascar, leading to the similar age estimates we recovered. Although our sampling is significantly improved relative to studies thus far, taxa from other geographic areas remain unsampled, and therefore our age and biogeographic estimates may change with an even more comprehensive dataset.

#### 4.2.1 The Voyage of *Ledebouria* Out of Africa, or *Ledebouria* Clade B

Reconstructions place the ancestral area of *Ledebouria* Clade B in the Zambezian region followed by migration into other regions, such as Yemen and India, between ∼20–15 mya (Figures 2 and 3). Although our age estimates do not allow us to unequivocally favor vicariance over dispersal from Africa to Eurasia, the current distribution of the Ledebouriinae, coupled with our knowledge of their biology and the overall results from this study, do allow us to form hypotheses of how the group may have migrated out of Africa. The Arabian Peninsula and Socotra were connected or in close proximity with mainland Africa from at least ∼17 mya (Bottenberg, 2012; Culek, 2013; Edgell, 2006; Fleitmann et al., 2004; Jacobs, 2004; Pirouz et al., 2017; Rögl, 1999), which would have allowed overland dispersal between these regions. Whether arrival into the Arabian Peninsula occurred before the full formation of the Red Sea or through the persistent connection offered by the Sinai Peninsula, will always be hard to assess with any confidence. Later, migration into Eurasia may have been facilitated by the *Gomphotherium* land bridge, which was present between ∼19–13 mya (Bialik et al., 2019; Okay et al., 2010; Rögl, 1999; Sen, 2013). The node reconstructed as India + Zambezian is estimated at 15.6 myr (95% CI 16.3–13.1), and therefore, we hypothesize the land bridge may have allowed dispersal of Ledebouriinae into Eurasia, a hypothesis also inferred for several animal lineages (Harzhauser et al., 2007; Sen, 2013). Additionally, as the Socotra archipelago drifted away from the mainland, we also hypothesize that vicariance likely promoted allopatric speciation in the region, which is evidenced by the different species found on different islands (Miller & Morris, 2004). Our age estimates, however, cannot fully rule out multiple dispersals to the archipelago, but the many species found within the island chain led us to favor a vicariant origin for these taxa. The vicariance hypotheses put forth here are not unique and fit a pattern recovered in multiple studies. In other groups with widespread distributions between Africa, the Arabian Peninsula, Socotra, and/or southern Asia, such as *Isodon* (Lamiaceae) (Yu et al., 2014), *Aganope* (Fabaceae) (Sirichamorn et al., 2014), *Searsia* (Anacardiaceae) (Yang et al., 2016), *Smilax* (Smilacaceae) (Chen et al., 2014), and several reptiles (Smíd et al., 2013; Tamar et al., 2016), vicariance has been hypothesized or alluded to for explaining distributions and biogeography, and many of these lineages have similar age estimates and distributions as *Ledebouria* Clade B. Lastly, pollen and wood fossils indicate historical widespread distributions between eastern Africa and south Asia of closely related taxa (Bonnefille, 2010; Morgan et al., 1994), and fossils indicate that forested habitats were present in the Arabian Peninsula during the Eocene and Oligocene, that later gave way to more open, xeric, grassland habitats in the Miocene (Jacobs et al., 1999; Whybrow & McClure, 1980). The modern-day equivalents of these conditions are commonly inhabited by Ledebouriinae (Lebatha, 2004; Venter, 1993). However, given phylogenetic and divergence times uncertainty and limited taxon sampling from this region, we cannot yet completely rule out multiple dispersals to the Arabian Peninsula and/or the Socotra Archipelago. Regardless, in summary, our age estimates and biogeographical reconstructions as well as the limited dispersal ability of Ledebouriinae, lead us to hypothesize that *Ledebouria* Clade B dispersed out of Africa to Eurasia via overland migration. The historical, widespread distribution was subsequently fragmented by Miocene-driven aridification.

#### 4.2.2 Within Africa—*Ledebouria* Clade A

Our results suggest that *Ledebouria* Clade A originated slightly later and in a more southerly region than that of *Ledebouria* Clade B (Table 1; Figures 2 and 3). Based on our dataset, *Ledebouria* Clade A arose ∼25–23 mya and remained predominantly within sub-equatorial Africa (Figures 2 and 3). Despite the low resolution within some areas of Clade A (Howard et al., 2022), we recovered multiple subclades that reflect geographic affinity (Figure 2). For example, we see a subclade of mixed geographic composition (i.e., *L. caesiomontana* + *L*. sp. 12 CCH190) (node c, Figure 2) that suggests a widespread distribution between the Kalahari and Zambezian regions that diverged and lead to *in situ* speciation in the Zambezian region followed by a dispersal to Madagascar (Figure 3). We recovered an additional subclade of predominantly Zambezian taxa (i.e., *L*. sp. 37 CCH169 + *L*. sp. 24 Uganda) (subclade in node a, Figure 2) with evidence of a dispersal to the Sudanian region (i.e., *L*. sp. 24 Uganda). Furthermore, two subclades of predominantly Namibian taxa (i.e., *L*. sp. 14 CCH109 + *L*. sp. 18 CCH218 (node b, Figure 2), and *L*. sp. 7 CCH066 + *L*. sp. 32 CCH145 (subclade in node a, Figure 2) suggest multiple dispersals to the region followed by *in situ* speciation (Figure 3). Lastly, a subclade of mostly South African taxa (i.e., *L. galpinii* + *L. coriacea*) (subclade in node a, Figure 2) was recovered (Figure 3). Overall, our results suggest that within a relatively short time frame (between ∼20–15 mya), a complex history of dispersals followed by *in situ* speciation events occurred within *Ledebouria* Clade A in southern Africa (Figure 3; Table S2).

From the Miocene onwards, complex interactions between mountain uplift and climate change promoted diversification across lineages within southern Africa (García-Aloy et al., 2017; Maswanganye et al., 2017; Neumann & Bamford, 2015; Nielsen et al., 2018). *Ledebouria* Clade A may have originated in southeastern Africa then dispersed and diversified as the region grew more heterogeneous and arid. For example, we found an instance of potentially rapid radiation within *Ledebouria* Clade A between ∼20–17 mya (Figure 3b), which corresponds to a clade with consistently poor phylogenetic resolution (Howard et al., 2022). This subclade contains taxa from across southern sub-Saharan Africa with geographic signals within (i.e., clades of South African, Namibian, and Zambian taxa) (Figures 2 and 3). It is possible that the Middle Miocene Climatic Optimum or Middle Miocene Climate Transition (Zachos et al., 2001) coupled with renewed uplift of the Great Escarpment spurred this radiation, and we see a steady rise in lineage accumulation from ∼20–15 mya (Figure 3b). During the Middle Miocene, the vegetation of South Africa was likely tropical, which gave way to more xerophytic shrubland by Late Miocene due to a global cooling climate (Pound et al., 2012). This time period is associated with diversification in the Crassulaceae (Lu et al., 2022) and also the expansion of C4 grasslands/savannas at a global scale (Bobe, 2006; Senut et al., 2009). Current biogeographical patterns of *Aloe* suggest dispersal from southern Africa to East Africa as seasonality and aridification increased across the continent within the past ∼16 myr (Grace et al., 2015). *Monsonia* (Geraniaceae) share similar distributional patterns as *Ledebouria* Clade A with multiple lineages comprised of mainly southwestern or southeastern taxa (i.e., southern African split between eastern and western clades), which radiated within the past ∼20–15 myr (García-Aloy et al., 2017). Overall, we find a fascinating, albeit more complicated, evolutionary history of *Ledebouria* Clade A within southern sub-Saharan Africa that remains to be fully resolved.

### 4.3 Limited, but intriguing insights into *Drimiopsis*

Based on our limited sampling of *Drimiopsis*, we find additional evidence for a southern and northern sub-Saharan African divide (Figures 2 and 3), which is also seen in ungulates (Lorenzen et al., 2012), *Lycaon pictus* (Canidae) (Marsden et al., 2012), *Encephalartos* (Cycadaceae) (Mankga et al., 2020), and *Agama* (Reptilia) (Leaché et al., 2014). The eastern/northern sub-Saharan African *Drimiopsis* arose ∼19–17 mya (Figure 2), during a time of increased mountain building in the region (∼17 and ∼13.5 mya; Wichura et al., 2015) that was followed by a shift towards a more arid climate (Bonnefille, 2010; Linder, 2017; Morgan et al., 1994; Senut et al., 2009). Several East African *Drimiopsis* have overall more succulent leaves compared to their southern African relatives (Lebatha, 2004), which may have been selected for as eastern Africa experienced greater aridification compared to the southeast. Additionally, East African *Drimiopsis* are known polyploids (Lebatha, 2004; Stedje & Nordal, 1987), which may have further spurred evolution in this region since polyploidy may provide an advantage in particular environmental conditions (e.g., aridity) (Manzaneda et al., 2012; Sonnleitner et al., 2010). In contrast, South African *Drimiopsis* occur in moist and shady habitats, and are not found in the drier portions of southwestern Africa (Lebatha, 2004). Given the estimated age of South African *Drimiopsis* (∼13.6 mya) (Figure 2), uplift of the Great Escarpment may have increased aridity in southwest Africa (Bobe, 2006; Partridge & Maud, 1987) that hindered *Drimiopsis* establishment in this region, but at the same time promoted diversification in southeastern Africa leading to its high diversity in the area (Lebatha, 2004). However, far greater sampling of *Drimiopsis* from its entire distribution is needed to test hypotheses regarding their evolutionary history. Ideally, an expanded phylogenetic framework would be coupled with ploidal inferences, anatomical studies, and experimental manipulations to fully assess the relationship between historical biogeography and morphological evolution.

### 4.4 Where to sample next?

Our taxon sampling heavily consisted of southern sub-Saharan African taxa (Figure 1), and largely excluded *Resnova*. Therefore, increased sampling of *Resnova* as well as *Ledebouria* and *Drimiopsis* from western, central, and northern Africa will provide further insights into the biogeographical history of the Ledebouriinae. For example, *Ledebouria* occurs in northern Angola (Gregory Jongsma, pers. obs.) and Gabon (Figure 1). This leads us to wonder whether a continuous distribution was once present along the central/western Atlantic coast of Africa, or if the Central African tropical forests have consistently obstructed Ledebouriinae dispersal within the coastal, wet tropics. Furthermore, a *Ledebouria* accession from Uganda is nested within *Ledebouria* Clade A, which suggests that by increasing sampling from northern/western Africa we may uncover additional biogeographic events in this clade. Lastly, multiple, morphologically distinct *Ledebouria* taxa are recorded from Socotra and surrounding islands (Miller & Alexander, 2010; Miller & Morris, 2004), and India (Giranje & Nandikar, 2016). Incorporating these into future phylogenetic studies may highlight further dispersals, or bolster support for allopatric speciation/vicariance as a major evolutionary process leading to the current distribution of the group outside of Africa.

## 5 CONCLUSION

The Ledebouriinae are a widespread, bulbous lineage of monocots that remain poorly understood taxonomically due to a high degree of phenotypic plasticity and significant under collection of specimens. Our study provides a pivotal first step towards refining our understanding of historical evolution and biodiversity within this fascinating lineage. The results of our analyses highlight the complex biogeographic history of the Ledebouriinae within and outside of sub-Saharan Africa. We find that the Ledebouriinae evolved within the past ∼30 myr, which suggests the group radiated in response to increasing climatic seasonality and orogenic activity since. Overall, given the evidence, we hypothesize that vicariance led to the current distribution of *Ledebouria* in Eurasia, with overland dispersal out of Africa potentially occurring via the *Gomphotherium* land bridge. However, long distance dispersal to Socotra and southern Asia cannot be fully ruled out at this time. We also find evidence of two independent dispersals to Madagascar. Within Africa, we can infer a northern and southern sub-Saharan division between the two *Ledebouria* lineages and within *Drimiopsis*. We also uncover a complex history of dispersals and *in situ* speciation events within southern Africa in *Ledebouria* Clade A, a clade that warrants improved phylogenetic resolution to further elucidate the processes affecting its current distribution. In conclusion, our study shows the value of increasing research focus on understudied lineages inhabiting seasonal and arid habitats.

## Acknowledgements

We are extremely grateful to Silke Rügheimer and Esmerialda Strauss at the National Botanical Research Institute in Windhoek, Namibia for their assistance with fieldwork and specimen export while in Namibia. We would also like to thank Inge Pehlemann for being a wonderful companion during excursions in search of Namibian *Ledebouria*. We are grateful to the governments of Namibia, Zambia, and Tanzania for issuing collection and export permits. Namibian collections were made under permit numbers 1784/2013, 1908/2014, 2056/2016, and 2185/2016. Collections from Zambia were made under permit number TJ/DNPW/101/13/18. Tanzanian collections were made under permit number 2017-22-NA-2016-247. Plants were imported under USDA permit numbers P37-09-00910, P37-16-00181, and P37-16-01462. Many thanks to Dylan Hannon, Gottfriend Milkuhn, Tom Cole, and Tom McCoy for providing leaf material from their collections. Lastly, we thank Killian Fleurial, Taylor LaVal, and Emily B. Sessa for assistance while in the field, and William Baker for assistance in obtaining sequence data for outgroup taxa. Funds from the Huntington Botanical Gardens, the Cactus and Succulent Society of America, the Pacific Bulb Society, the San Gabriel Cactus and Succulent Society, the Florida Museum of Natural History, the American Society of Plant Taxonomists, the University of Florida International Center, the University of Florida Department of Biology, the Botanical Society of America, the Society of Systematic Biologists, Xeric Growers, and numerous private donors helped to support this work.

## Data Availability

Raw reads from PAFTOL are available in ENA (accession PRJEB35285) and assembled reads from www.treeoflife.kew.org. Raw reads of the Ledebouriinae are available from the SRA (accession PRJNA721471). Alignments, species trees, biogeography inputs/outputs are all available on Dryad (FOR REVIEW ONLY: https://datadryad.org/stash/share/ApGYMXXwCCd1G873IOkciaD27YdQyNqg0lpvKT38Xic).

## Biosketch

Cody Coyotee Howard is an Assistant Professor at Oklahoma State University broadly interested in geophyte evolution and biogeography, and he has a particular fascination with bulbous plants. The Ledebouriinae represent one bulbous lineage that has captivated him for years, and these plants will continue to do so for many more.

## Author contributions

Conceptualization CCH, TSH, NC; Formal analysis, CCH and ARZ; Investigation, CCH and ARZ; Resources, TSH, LN, DC, LN, PM, ARZ, NC; Writing — Original Draft, CCH and NC; Writing — Review & Editing, all authors; Visualization, CCH; Funding acquisition, CCH and NC.

## Supporting information

**Table S1.** Fossil and secondary calibration points used in the penalized likelihood (treePL) analysis. With the exception of monocots, all calibrations were set as minimum values in the analysis.

| node | minimum calibration | source |
| --- | --- | --- |
| Zingiberaceae | 72.1 | Iles et al. (2015) |
| Typhaceae | 51.66 | Iles et al. (2015) |
| Cyperaceae | 47.0 | Iles et al. (2015) |
| Amaryllidaceae | 48.88 | Pigg et al. (2018) |
| Coryphoideae | 83.6 | Iles et al. (2015) |
| Agavoideae | 14.5 | Iles et al. (2015) |
| Xanthorrhoeaceae | 38.0 | Iles et al. (2015) |
| Goodyerinae | 15.0 | Iles et al. (2015) |
| Monocots | 131–135 | Magallón et al. (2015) |

**Table S2.** Top ten most common transition types found in a subclade of *Ledebouria* Clade A (Figure 3b) using stochastic mapping of biogeographic events.

| starting_range | ending_range | average_n_events | SD | transition_type |
| --- | --- | --- | --- | --- |
| K | K | 23.432 | 2.121 | in situ speciation |
| Z | Z | 19.252 | 1.297 | in situ speciation |
| K | KN | 3.393 | 0.517 | dispersal |
| T | T | 2.764 | 2.089 | in situ speciation |
| C | C | 1.29 | 1.912 | in situ speciation |
| KZ | K | 1.09 | 0.307 | vicariance or subset sympatry or extinction |
| KZ | Z | 1.09 | 0.307 | vicariance or subset sympatry or extinction |
| K | KZ | 1.029 | 0.215 | dispersal |
| CK | C | 0.912 | 0.703 | vicariance or subset sympatry or extinction |

**Figure S1.** Full dated phylogeny with 95% confidence internals shown.

**Figure S2.** Biogeographical reconstruction using the DIVA-like model with nodes showing the probable range for each region at that node.

