## Supplementary figures and images for "From southern Africa and beyond: historical biogeography of a monocotyledonous bulbous geophyte"

### Figure S1

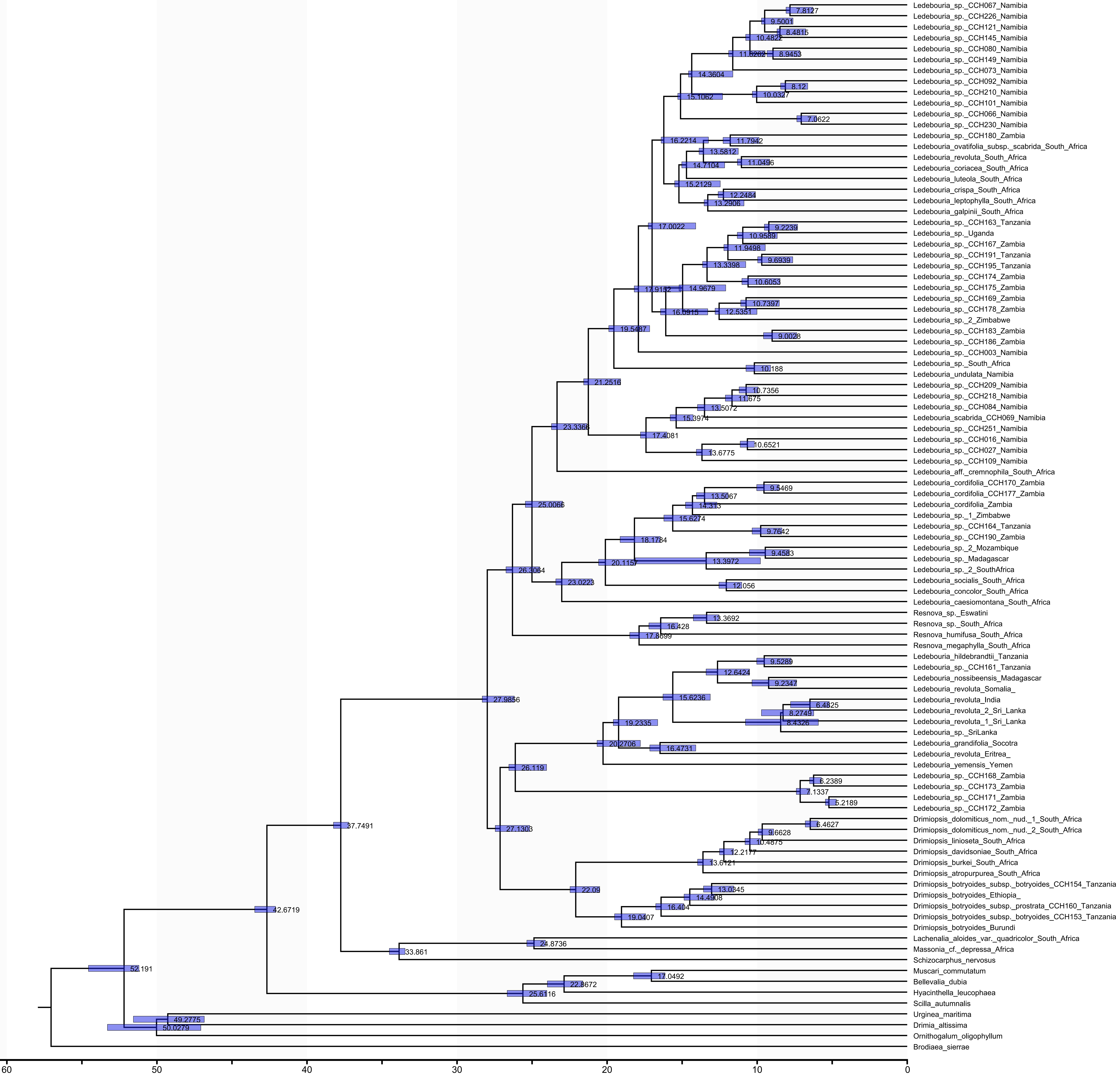

### Figure S2

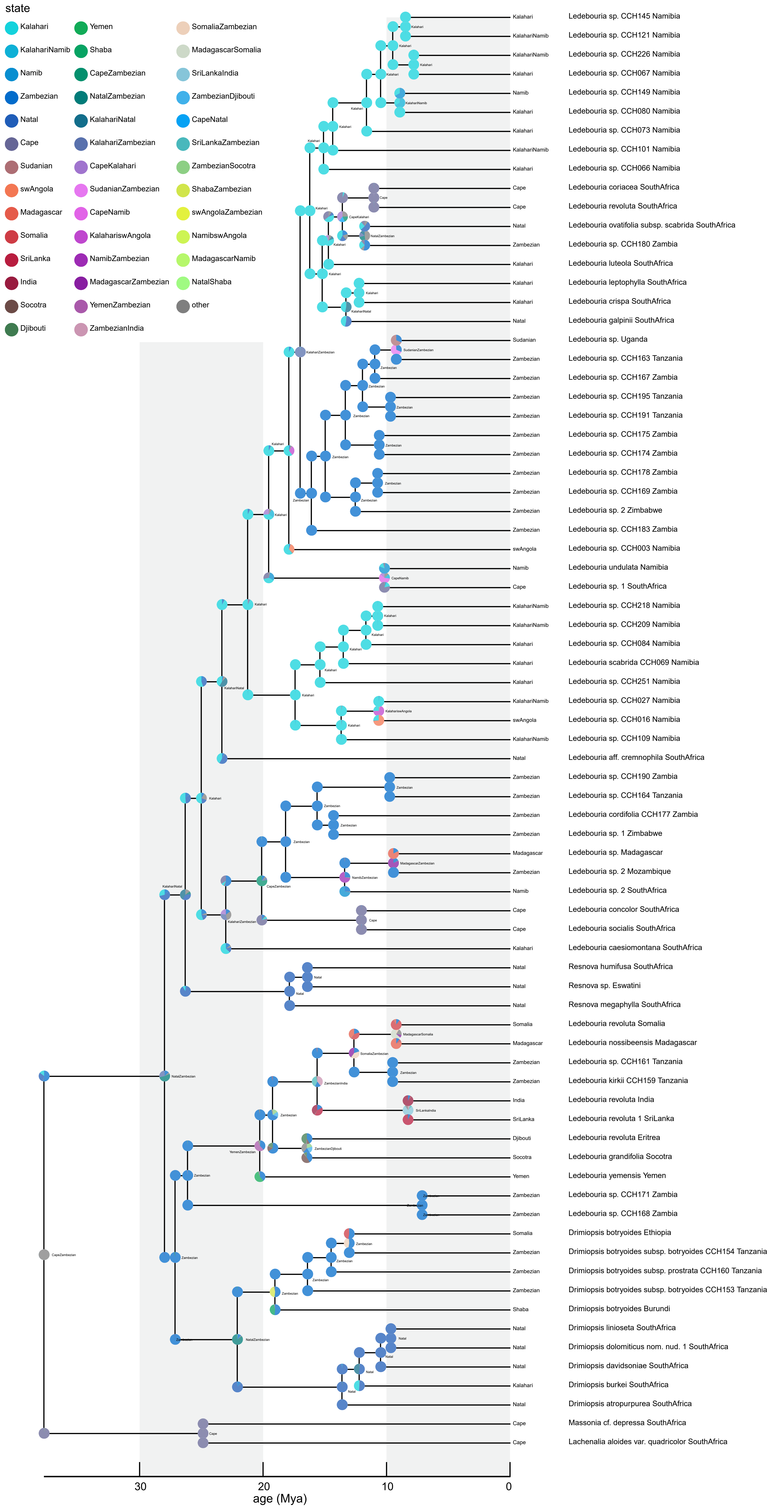
